# Speaker-induced suppression in EEG during a naturalistic reading and listening task

**DOI:** 10.1101/2022.12.13.519519

**Authors:** Garret L. Kurteff, Rosemary A. Lester-Smith, CCC-SLP, Amanda Martinez, Nicole Currens, Jade Holder, Cassandra Villarreal, Valerie R. Mercado, Christopher Truong, Claire Huber, Paranjaya Pokharel, Liberty S. Hamilton

**Author notes:** **Corresponding Author:** Liberty S. Hamilton.

## Abstract

Speaking elicits a suppressed neural response when compared to listening to others’ speech, a phenomenon known as speaker-induced suppression (SIS). Previous research has focused on investigating SIS at constrained levels of linguistic representation, such as the individual phoneme and word level. Here we present scalp EEG data from a dual speech perception and production task where participants read sentences aloud then listened to playback of themselves reading those sentences. Playback was separated into predictable repetition of the previous trial and unpredictable, randomized repetition of a former trial to investigate the role predictive processing plays in SIS. Concurrent EMG was recorded to control for movement artifact during speech production. In line with previous research, event-related potential analyses at the sentence level demonstrated suppression of early auditory components of the EEG for production compared to perception. To evaluate whether specific neural representations contribute to SIS (in contrast with a global gain change), we fit linear encoding models that predicted scalp EEG based on phonological features, EMG activity, and task condition. We found that phonological features were encoded similarly between production and perception. However, this similarity was only observed when controlling for movement by using the EMG response as an additional regressor. Our results suggest SIS is at the representational level a global gain change between perception and production, not the suppression of specific characteristics of the neural response. We also detail some important considerations when analyzing EEG during continuous speech production.

## Introduction

### Background

Speech production and speech perception are frequently studied separately in research, yet the two processes have a robust, interactive theoretical link (Houde and Nagarajan 2011; Tourville and Guenther 2011; Zheng, Munhall, and Johnsrude 2010; Watkins, Strafella, and Paus 2003; Skipper, Devlin, and Lametti 2017). Models of the neurobiology of speech production universally include the sensorimotor control of speech, a mechanism by which speakers can detect errors via auditory and somatosensory feedback and subsequently correct those errors (Perkell et al. 1997; Tourville and Guenther 2011; Parrell et al. 2019; Houde and Chang 2015). Errors are identified in part by comparison to the *efference copy*, which represents a set of sensory expectations about the content of an utterance that is generated during pre-articulatory planning of an utterance (Hawco et al. 2009; Hashimoto and Sakai 2003; Zheng, Munhall, and Johnsrude 2010; Behroozmand and Larson 2011; Greenlee et al. 2013). Difficulty with pre-articulatory planning of speech as well as disruption of the efference copy during speech production are theorized to be neurological mechanisms responsible for stuttering (Max and Daliri 2019; Toyomura et al. 2020; Smith and Weber 2017). Deficits in this mechanism have also been observed in people with schizophrenia (Heinks-Maldonado et al. 2007; McGuire et al. 1995; Woodruff et al. 1997) and Parkinson’s disease (Hoffman 2014).

When comparing speech production to perception, a phenomenon known as speaker-induced suppression (SIS) has been observed, where neural responses to (errorless) self-generated sounds are suppressed in relation to externally generated sounds (Martikainen, Kaneko, and Hari 2005; Brumberg and Pitt 2019; Niziolek, Nagarajan, and Houde 2013; Houde et al. 2002). The exact neural mechanisms behind SIS are not well understood, but many EEG (and MEG) studies point to early auditory components such as the N100(m) being a potential biomarker of the efference copy (Heinks-Maldonado et al. 2007; Martikainen et al. 2005; Behroozmand & Larson 2011). Work in animal models has suggested direct feedback from the motor cortex to inhibitory neurons in the primary auditory cortex suppress responses to self-generated sounds before and during movement (Schneider, Nelson, and Mooney 2014). The behavioral explanation for SIS is that it is responsible for distinguishing internally and externally generated speech for the purposes of speech motor control (Houde et al. 2002; Houde and Nagarajan 2011); however, a neurophysiological explanation for SIS is unclear. It is widely accepted that the brain uses some sort of intermediate representations when processing language from its constituent acoustic signal (Mesgarani et al. 2014; Appelbaum 1996), and specific representations of the perceptual response may be deemed unnecessary during production (e.g., phonological features, acoustic properties of the speech signal, etc.). The suppression of specific representations during speech production could explain the neurophysiological basis of SIS; alternatively, SIS could be explained as a general suppression of the whole neural response.

While speech perception does involve feedforward expectations (Poeppel and Monahan 2011), the predictability of speech perception is not as complete as speech production, due to expectations about utterance content being internally generated during utterance planning. Thus, stimulus predictability would appear to be a fundamental difference between conditions where speech is suppressed and where speech is not suppressed, and offers a potential explanation for the research question of *what* is being suppressed during SIS. Studies of altered auditory feedback, in which self-generated speech is acoustically perturbed in real time, show predictable patterns of feedback perturbation elicit larger corrective responses than unpredictable ones (Lester-Smith et al. 2020), which may corroborate the link between SIS and predictability.

Until recently, electroencephalography (EEG) studies of speech production were highly constrained in both the content of the produced speech and the analyses available to researchers due to challenges intrinsic to studying speech production that do not hinder the study of speech perception. For example, speech production studies were unable to advance beyond the single word level and frequently were epoched to events other than the onset of articulation (e.g., stimulus presentation, offset of articulation) in an effort to prevent electromyographic (EMG) artifact associated with articulation from contaminating the neural response (Shuster 2003; Okada, Matchin, and Hickok 2018; Singh et al. 2018; Jiang, Bian, and Tian 2019). Fortunately, advances in artifact correction techniques have resulted in the study of speech production via EEG above the word level. Ries et al. 2021 recently demonstrated the feasibility of analyzing EEG responses to multi-word production. Shifting EEG studies towards language as it occurs in natural settings – compared to the heavily constrained single word or syllable-level studies of the past – facilitates generalization to clinical applications and reinforces the interdisciplinary drive to use more ecologically valid stimuli in studies of the neural representation of speech and language (Hamilton and Huth 2020; Matusz et al. 2019). Studies which expand beyond using evoked stimuli and incorporate naturalistic stimuli (e.g. sentences) raise the ecological validity of the research while also providing a window of analysis for the feedforward and feedback processes that link perception and production (Casserly and Pisoni 2010; Houde and Nagarajan 2011; Poeppel and Monahan 2011; Kearney and Guenther 2019).

### Current Study

In this study, we aim to investigate differences in EEG responses between sentence-level speech perception and production, as well as predictable and unpredictable speech perception to discover what is driving the suppression observed during speech production. Is this suppression a universal feature of predictable auditory stimuli, or is it inherent to generating an utterance? Does phonological tuning to specific speech features (Di Liberto, O’Sullivan, and Lalor 2015; Desai et al. 2021; Khalighinejad, Cruzatto da Silva, and Mesgarani 2017) differ during production and perception? To investigate these questions, we designed an experiment that used identical acoustic stimuli in separate speech perception and production conditions then compared the difference in event-related potentials as well as in tuning of phonological features across conditions. We hypothesized that, although speech production will be suppressed relative to perception in this study, phonological feature tuning would remain stable between modalities of speech. Additionally, we expect a similar trend in unpredictable perceptual stimuli, such that representations will remain stable but show a general enhancement relative to predictable perceptual stimuli. Elucidating the neurophysiological explanations underlying the phenomenon of speaker-induced suppression has the potential to improve assessment and treatment of neuropsychological disorders in which this phenomenon is affected.

## Materials & Methods

### Participants

21 participants (11F, age 24.4±3.9) were recruited from The University of Texas at Austin. This is in line with sample sizes of recent EEG studies of speech production (Ries et al. 2021; Zhao and Rudzicz 2015; Goregliad Fjaellingsdal et al. 2020). All participants were native speakers of English with typical hearing as assessed through pure tone audiometry and a speech-in-noise hearing test (QuickSIN, Interacoustics). Participants provided written consent for participation in the study and were compensated at a rate of $15/hr with an average session length of 2 hours (1 hour for setup, 1 for recording EEG). One participant was excluded due to a recording error, leaving 20 participants in the final analysis. All experimental procedures were approved by the Institutional Review Board at The University of Texas at Austin.

### Materials

The task was designed using a dual perception-production block paradigm, where trials consisted of a dyad of sentence production followed by sentence perception. In each trial, participants overtly read a sentence, then listened to a recording of themselves reading the produced sentence. Perception trials were divided into blocks of predictable and unpredictable stimuli. *Predictable* stimuli consisted of immediate playback of the production trial, while *unpredictable* stimuli consisted of a randomly selected production trial from the previous block. A schematic is provided in Figure 1A. The generation of perception trials from the production aspect of the task allowed stimulus acoustics to be functionally identical across conditions. Sentences were taken from the MultiCHannel Articulatory (MOCHA) database, a corpus of 460 sentences that include a wide distribution of phonemes and phonological processes typically found in spoken English (Wrench 1999). These sentences have been used previously in intracranial studies of speech production (Chartier et al. 2018). A subset of 50 sentences (100 for the first two participants) from MOCHA were chosen at random for the stimuli in the present study; however, before random selection, 61 sentences were manually removed by an author (GLK) for either containing offensive semantic content or being difficult for an average reader to produce to reduce extraneous cognitive effects and error production, respectively. We changed the sentence set from 100 to 50 sentences after the first two participants due to concerns about participant fatigue during the task. Participants completed six blocks of the task for a total of 300 perception and 300 production trials per participant (400 for the first two participants). Sentences had a median length of 2.9 seconds. A broadband click tone was played in between trials as an additional cue to assess the effect of EMG correction on low level auditory responses.

**Figure 1.**
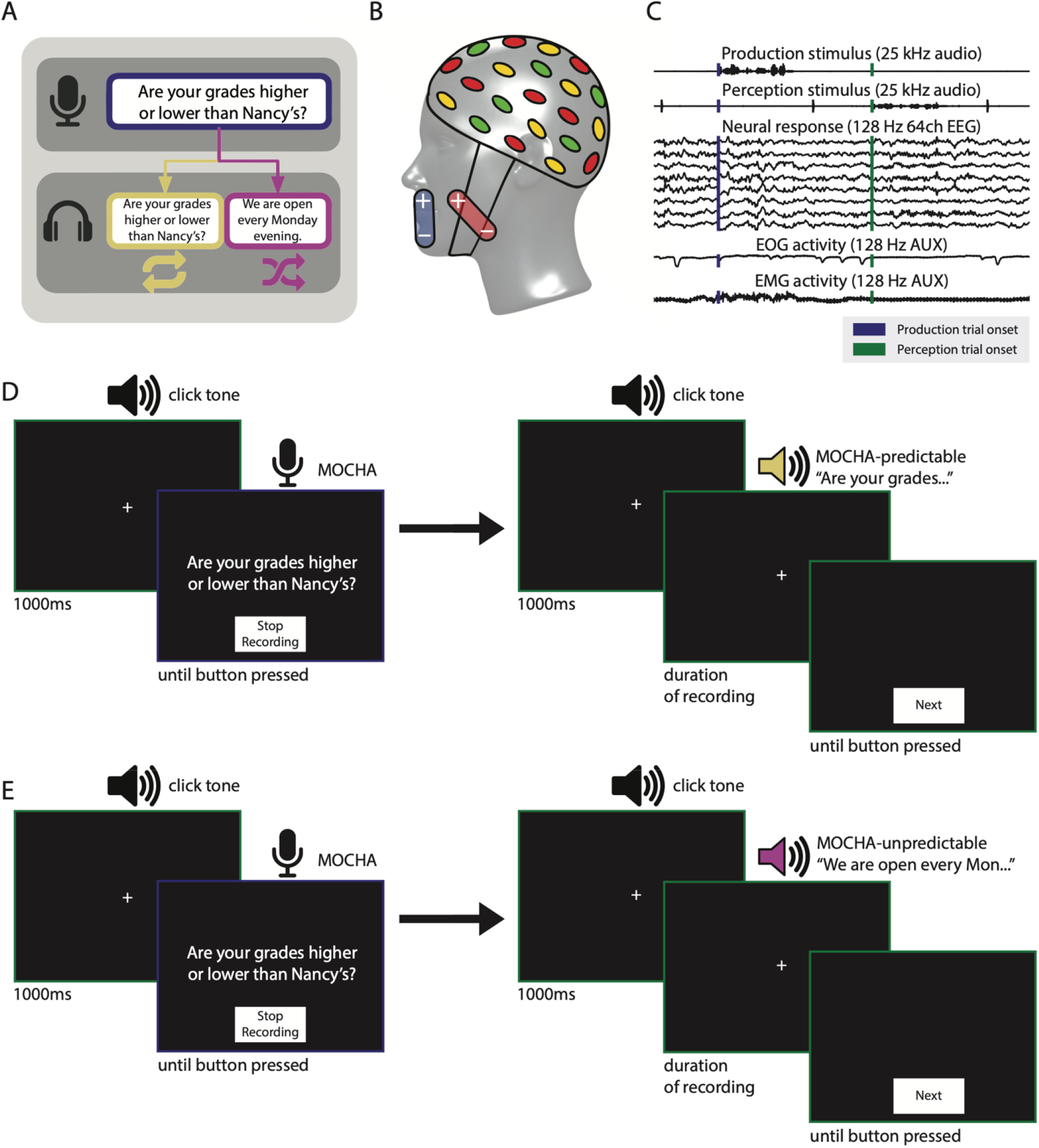
Dual perception-production task and EEG data collection schematic. A: Schematic of trial types in the task. The participant first reads a sentence aloud (indigo), then hears playback of the same audio (yellow, predictable condition) or audio from a different random trial (magenta, unpredictable). B: Schematic of auxiliary EMG electrode placement on orbicularis oris (blue) and masseter (red). C: Visualization of all signals recorded during task, including produced audio (speech), perceived audio (clicks and speech), and EOG and EMG channels. Only eight EEG channels are visualized here, but a total of 64 were recorded and used in analysis. Vertical lines denote the beginning of a production (indigo) or perception (green) trial. Blinks are observed as deflections in the EOG channel; muscle activation during production is notable as high activity in the EMG channel. D, E: Outline of trial procedure for predictable (yellow) and unpredictable (magenta) blocks.

Stimuli were presented in a dimly lit sound-attenuated booth on an Apple iPad Air 2 using custom interactive software developed in Swift (Apple XCode version 9.4.1). Auditory stimuli were presented at a comfortable listening level via foam-tipped insert earbuds (3M, E-A-Rtone Gold 10_Ω_, Minnesota, USA). Visual stimuli were presented in a white font on a black background after a 1000ms fixation cross to minimize visual artifact in the EEG signal (Figures 1D, 1E). Accurate stimulus presentation timing was controlled by synchronizing events to the refresh rate of the screen. The iPad was placed on a table over the participants’ lap so they could advance trials during the task with minimal arm movement. Participants were instructed to complete the task at a comfortable pace and were familiarized with the task before recording began. Trial information, including onset and offset of each trial, transcriptions of produced and heard sentences, trial type, trial number, and block number were collected by an automatically generated log file to assist in data processing.

### EEG Data Collection

64-channel scalp EEG and audio were recorded continuously via BrainVision actiChamp amplifier (Brain Products, Gilching, Germany) with active electrodes at 25kHz. A high sampling rate was used to synchronize task audio and EEG, which were recorded using the same amplifier. Conductive gel (SuperVisc, EASYCAP) was applied to the scalp at each electrode, and impedance at each electrode was kept below 15kΩ throughout the recording. Audio signals from both the insert earphones (presented audio) and microphone (produced audio) were captured as additional EEG auxiliary channels and were aligned with neural data via a StimTrak processor (Brain Products). Vertical electrooculography (vEOG) was captured via auxiliary electrodes above and below the left eye in line with the pupil. Auxiliary electrodes were also used to capture facial EMG activity (Figure 1B); these electrodes were placed on the orbicularis oris and mandible in the majority of participants (N=11), but on other muscles important to articulation (masseter (N=6), submental triangle (N=2)) in several participants (Stepp 2012; Van Eijden, Blanksma, and Brugman 1993; Rastatter and De Jarnette 1984). Multiple placements were utilized due to issues with electrode adherence caused by participant facial hair. All placements were trialed on a participant who consented to additional time during setup. A reference electrode for all auxiliary electrodes was placed on the left earlobe. Auxiliary EMG placement was not required for preprocessing but provided validation that EMG artifact was removed during preprocessing. EMG activity associated with the onset of articulation, which caused the largest artifacts in the temporal window of interest for event-related potential analysis of speech production, was automatically detected and epoched from auxiliary EMG channel activity. All recorded signals timed according to stimulus onsets are visualized in Figure 1C. The first two participants did not have auxiliary electrode placement due to unavailability of recording hardware, so EMG activity was corrected based on EEG channels only (see below).

### EEG Data Processing

All EEG processing was performed offline using custom Python scripts and functions from the MNE-python software package (Gramfort et al. 2014). EEG, EOG, and EMG data were downsampled from 25kHz to 128Hz prior to analysis. EEG data were referenced to the linked mastoid electrodes (the average of the TP9 and TP10 channels) and notch filtered at 60Hz to remove line noise. For one subject (OP17), one reference electrode was a bad channel and was interpolated prior to re-referencing. The data were next filtered from 1-30Hz (Hamming window, 0.0194 passband ripple with 53 dB stopband attenuation, 6 dB/octave). Bad channels and segments were manually rejected, then Independent Component Analysis (ICA) was performed to correct for EOG and electrocardiographic (EKG) artifact with the number of components equal to the number of good channels. ICA components related to vEOG, horizontal EOG (hEOG) and EKG were manually identified and removed via scalp topography and epoching component activity to vEOG activity (obtained via MNE function create_eog_epochs). The selected ICA components were next removed from the unfiltered data. After ICA, data were filtered at 0.16Hz and corrected for EMG artifact via blind source separation algorithm based on Canonical Correlation Analysis (CCA, (De Clercq et al. 2006)), a technique that has been previously demonstrated to correct for EMG artifact in speech production EEG tasks (Vos et al. 2010; Ries et al. 2021; Riès et al. 2013). In line with these studies, CCA was performed in two passes: first, a 30-second window to remove tonic muscle activity; second, a 2-second window to remove rapid bursts of EMG associated with speech production. CCA was performed using the Automatic Artifact Removal plugin for EEGLab (Gómez-Herrero 2007). After CCA and prior to analysis, bad channels were interpolated and data were bandpass filtered between 1-30Hz.

### Analysis

#### Event-Related Potential Analysis

Accurate timing information for words, phonemes, and sentences was generated to allow epoching of EEG data to multiple levels of linguistic representation. A modified version of the Penn Phonetics Forced Aligner (Yuan and Liberman 2008) was used to automatically generate Praat TextGrids (Boersma and Weenink 2013) using a transcript generated by the iPad’s log file. Automatically generated TextGrids were checked for accuracy by authors AM, NC, JH, CV, VM, CT, CH, and PP. The first author (GLK) supervised the transcription process and checked the final TextGrids for accuracy before generating event files used in the analyses. Event files containing start and stop times for each phoneme, word and sentence, as well as information about trial type (perception vs. production), were created using the log files and TextGrids. A second set of event files corresponding to the inter-trial click sound were generated via a match filter process where the audio signal of the click was convolved with the EEG audio signal to find exact timing matches (Turin 1960).

To examine the differences between perception and production at the sentence level, sentence-level event files were used to epoch the neural response between -1.5s to +3s relative to sentence onset (Ozker et al. 2022). Epochs ±10 SD from the within-subject mean were rejected. Linear mixed-effects (LME) models were created and assessed using the lmertest package (Kuznetsova, Brockhoff, and Christensen 2017) in R to determine statistical differences between different task conditions within relevant time windows, specifically the N100 (80-150ms) and P200 (150-250ms). The peak amplitudes and latencies of these windows, as well as the peak-to-peak amplitude of the N100 and P200 components, were used as response variables. Latency was calculated as the time at which the largest peak within a time window of interest occurred. LME models were specified using the equation:

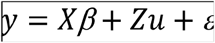

Where β represents fixed-effects parameters, *u* represents random effects, and *ε* represents residual error. *X* and *Z* are matrices of shape (*n* × *p*), where *n* is the number of observations of each parameter and *p* is the value of the parameter at observation *n*. In all models, the fixed effect was the response variable of interest (i.e., N100 & P200 amplitude & latency; peak-to-peak amplitude) and Subject was used as a random effect. F tests were calculated using Kenward-Roger approximation with *n* degrees of freedom specified (Kenward and Roger 1997).

### Linear Encoding Model Analysis

Linear encoding models (also referred to as spectrotemporal/multivariate temporal receptive field (s/mTRF) models in previous literature) were fit to describe the selectivity of the EEG responses to phonological features corresponding to place and manner of articulation (Crosse et al. 2016; Di Liberto, O’Sullivan, and Lalor 2015; Hamilton, Edwards, and Chang 2018; Mesgarani et al. 2014; Desai et al. 2021). This model takes the form of the equation below:

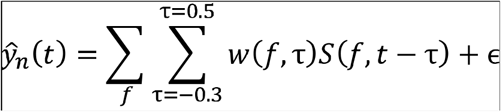

Where 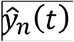 represents the estimated EEG signal for electrode *n* at time *t*. The stimulus matrix *S* consists of behavioral information regarding features (*f*) for each time point 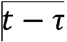, where 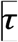 is the time delay between the stimulus and neural activity in seconds. Features included combinations of binary features for perception, production, predictable, and unpredictable trials, as well as continuous, normalized EMG activity recorded from auxiliary electrodes, and binary features for the presence of phonological features at each time point (as in (Desai et al. 2021; Hamilton, Edwards, and Chang 2018; Mesgarani et al. 2014). The “full” model stimulus matrix contained 14 phonological features as well as four binary features encoding trial information (perception, production, predictable, unpredictable) and normalized EMG activity from facial electrodes for a total of 19 features. These phonological features for place and manner of articulation were identical to those used in previous work (Desai et al. 2021; Hamilton et al. 2021; Mesgarani et al. 2014) and included sonorant, obstruent, voiced, nasal, syllabic, fricative, plosive, back, low, front, high, labial, coronal, and dorsal. Phonemes were coded in a binary matrix where a 1 indicated the onset of a phoneme’s articulation via timing information obtained from the TextGrids.

We fit separate models to predict the EEG response in each channel using time delays of -0.3s to +0.5s. This delay range encompassed the temporal integration times to similar responses found in previous research (Hamilton, Edwards, and Chang 2018) but with an added negative delay to encompass potential pre-articulatory neural activity (Chartier et al. 2018). Data were split 80-20 into training and validation sets. To avoid overfitting, the data were segmented along sentence boundaries, such that the training and validation sets would not contain information from the same sentence. These segments were then randomly combined into the 80/20 training/validation sets. Weights for each feature and time delay 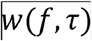 were fit using ridge regression on the training set and a regularization parameter chosen by 10 bootstrap iterations, fitting on subsets of the training set. The ridge parameter was selected at the value that provided the highest average correlation performance across all bootstraps. Ridge parameters between 10^−5^ and 10^5^ were tested in 20 logarithmically scaled intervals. Model performance was assessed using correlations between the EEG response predicted by the model and the true EEG response. Significance of these correlations was obtained through a bootstrap procedure with 100 iterations in which the training data were shuffled in chunks to remove the relationship between the stimulus and response but preserve temporal correlations within the EEG signal. Visual inspection of the data revealed two subjects (OP4, OP12) for whom responses showed no discernible receptive field structure even after greatly expanding the range of ridge parameters, motivating their exclusion from the analysis.

## Results

Topographic inspection of sentence-level ERP activity revealed a frontocentral ROI of nine channels that elicited the strongest response to sentence onset during speech perception and production (F1, Fz, F2, FC1, FCz, FC2, C1, Cz and C2). This ROI is used in the ERP results, but linear encoding models were fit on all channels for all subjects.

### Event-Related Potential Results

After verifying the integrity of the dataset, we wished to understand whether and how responses to continuous speech differ for production versus perception and for the predictable and unpredictable conditions. Sentence-level ERPs for both perception and production were epoched to the onset of sentence articulation (the first phoneme in the trial sentence). These ERPs demonstrated a relative suppression of EEG activity in production trials compared to perception trials (Figure 2B). The N1 and P2 components are present at the sentence level in both perception and production conditions but reduced in amplitude for the production trials. We fit LME models comparing perception and production in windows of interest(windowed amplitude ∼ Condition + (1|Subject) and windowed latency ∼ Condition + (1|Subject)). We found significantly lower amplitudes for N100 (Estimated Marginal Mean_perception-production_ = - 2.31±.15μV; *p*<.001) and significantly higher amplitudes for P200 (EMM_perception-production_ = 1.72±0.15μV; *p*<.001) during perception compared to production. This was also in line with increased peak-to-peak amplitude (EMM_perception-production_ = 3.96±0.15μV; *p*<.001) in perception compared to production. In addition, N100 latency was decreased in production compared to perception (EMM_perception-production_ = 1.6±0.4ms; *p*<.001), and similar results were seen for P200 latency (EMM_perception-production_ = 2.75±0.6ms; *p*<.001). Suppression during speech production relative to perception in this task highlights differences in processing internally and externally generated speech. A potential explanation for the suppression observed during speech production is that speech production contains expectations about the content of the utterance via efference copy, while speech perception requires information about the content of the utterance to be processed in real time.

**Figure 2.**
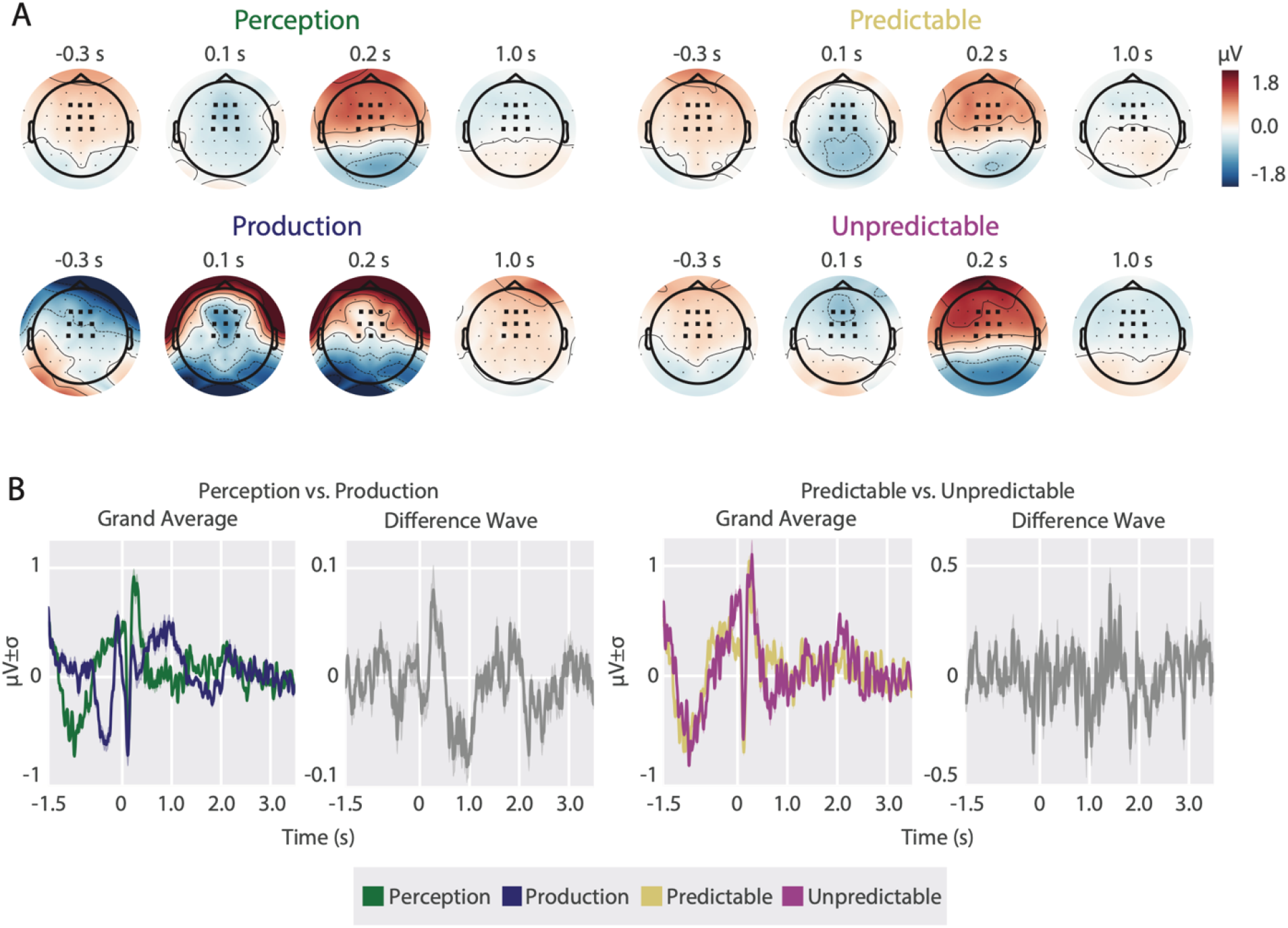
Event-related responses to sentence onset demonstrate suppression of N1-P2 during speech production. Speech production (indigo) is suppressed relative to perception (green), but no such difference is observable for predictable (yellow) vs. unpredictable (pink) speech perception. A: Topographic distribution of activity relevant to sentence onset for each task condition. Nine electrode ROI used in analyses represented via bolded electrodes. B: Grand average ERPs and difference waves comparing speech production and speech perception (left) and predictable and unpredictable speech perception (right).

Although differences were significant between perception and production trials, the differences between predictable and unpredictable speech perception were less pronounced: LME modeling (Window ∼ Condition + (1|Subject)) did not reveal a significant difference in N100 (EMM_predictable-unpredictable_ = 0.3±0.2μV; *p*=0.12) and P200 (EMM_predictable-unpredictable_ = - 0.2±.2μV; *p*=0.37) amplitudes across this contrast. However, peak-to-peak amplitude (EMM_predictable-unpredictable_ = -0.5±0.2μV; *p*=0.03) and N100 latency (EMM_predictable-unpredictable_ = 1.5±0.7ms; *p*=0.02) differed significantly between predictable and unpredictable trials, with an earlier response to unpredictable compared to predictable sentences. P200 latency did not differ significantly (EMM_predictable-unpredictable_ = 0.2±.1; *p*=0.84). Because predictable and unpredictable perception trials were split into blocks during the task, an oddball response was not elicited for the unpredictable stimuli. To further investigate the significance of peak-to-peak amplitude and N100 latency between predictable and unpredictable stimuli, a series of Wilcoxon signed-rank tests with Benjamini-Yekutieli correction (Benjamini and Yekutieli 2001) comparing N100-P200 peak-to-peak amplitude and N100 latency on a within-subject basis were performed. These significance tests revealed only three individual subjects that demonstrated a significant suppression between predictable and unpredictable speech perception (OP01 *p*=0.02; OP07 *p*<.001; OP21 *p*=0.0002), and only two subjects with a significant difference in N100 latencies (OP1 *p*=0.004; OP19 *p*=0.04). This within-subject analysis suggests the significance of peak-to-peak amplitude and N100 latency observed in the LME results is caused by outlier subjects rather than a generalizable effect. Overall, differences in predictability were less pronounced than the differences between perception and production trials. These minor differences between expected and unexpected speech perception suggest the suppression seen during speech production is not fundamentally linked to the predictable nature of speech production. In other words, feedforward processing of speech perception and feedforward processing of speech production reflect different neural mechanisms.

### Linear Encoding Model results

While our ERP results provide insight into the timing and magnitude of differences in responses during perception and production, they do not provide information regarding any potential differences in responses to specific speech features or content. Furthermore, ERP analyses are constrained by the need to average many trials that are time-locked to a particular event (Luck 2014). Thus, ERP analyses may not be as sensitive to uncovering differences outside of the onset of the sentence, or for specific phonological features within continuous speech. To address this limitation, we performed additional analyses where we fit linear encoding models for continuous production and perception. These analyses are powerful in that they allow for investigation of continuous, natural speech without the need for trial averaging. They also allow us to further probe specific differences (or lack thereof) in tuning across our different task conditions.

Model performance was evaluated by calculating the linear correlation coefficients (*r*) between the EEG response predicted by the model and the actual response. We also probed the importance of individual features on model performance by ablating specific features from the stimulus matrix *S* and observing the change in correlation coefficients between ablated and full models. Such variance partitioning methods have been used to uncover the unique variance explained by particular features (de Heer et al. 2017; Hamilton et al. 2021). For example, if a model that omitted normalized EMG predicted the neural response less accurately, the interpretation is that EMG contains important information for accurately modeling EEG activity. For each task-related feature in the “full” model (14 phonological + 4 task features; Figure 3C), we fit a separate model omitting that feature. Lastly, one model had two additional sets of phonological features (i.e., 14 phonological features during production + 14 phonological features during perception + 14 phonological features in either condition + 4 task features; Figure 3A). These were split by modality to observe if phonological feature tuning changed between perception and production. We call this model the “task-specific” encoding model, which is in comparison to the “identical” encoding model in which phonological feature tuning is assumed to be the same across all conditions, with only a baseline change fit by the condition features. The estimated marginal mean correlation coefficients of these models were compared via LME modeling with subject and channel location as random effects (r ∼ Model + (1|Subject) + (1|Channel)). Separating phonological tuning by the modality of speech (i.e., perception vs. production) had a significant effect on model performance (EMM_*r*(identical-separate)_ = - 0.012±0.002; *p*<.001), such that separating phonological feature tuning during production from phonological feature tuning during perception improved the model’s ability to predict the held-out neural response (Figure 3B). This result, which was contrary to our initial hypothesis, suggested that phonological feature encoding differs during speech perception and production. However, due to the influence of electromyographic artifact during speech production, speech perception in this task is a combination of sensory and motor responses, while speech perception in this task is purely sensory, which may explain the difference in the models presented in Figure 3.

**Figure 3.**
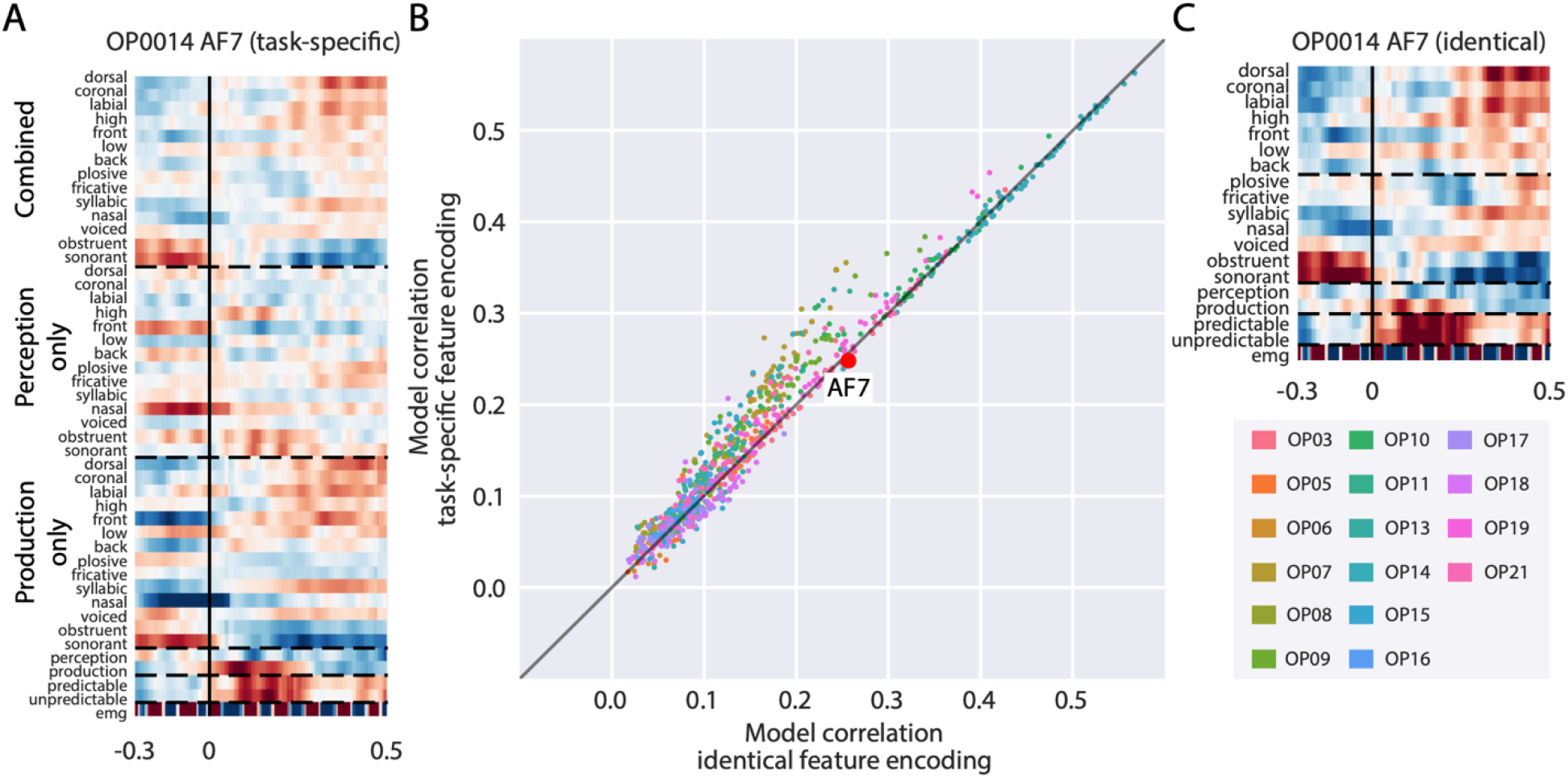
Separating phonological feature encoding by modality of speech improves model performance. A: Temporal receptive field for an individual electrode with stimulus characteristics divided by task condition (i.e., perception vs. production). B: Scatterplot of channel-by-channel correlation coefficients between two compared models, colored by individual subject. Diagonal black line represents unity. C: Temporal receptive field for an individual electrode with stimulus characteristics identical across task condition.

While we utilized methods to correct for EMG artifact that have been previously demonstrated in the literature to be successful (Ries et al. 2021; Vos et al. 2010; Chen et al. 2019), there is no definitive way to rule out residual EMG given the lack of ground truth in the sources that contribute to the electroencephalogram. As a result, we further explored the influence of EMG artifact on model performance by fitting linear encoding models that included normalized EMG activity recorded from auxiliary facial electrodes in tandem with the EEG as a regressor. Models that include or exclude the auxiliary EMG but are otherwise identical in their stimulus matrices were compared in an ablation-based approach to explore the contribution of specific features to model performance (Ivanova, Hewitt, and Zaslavsky 2021). Linear correlation coefficients were compared using an LME model identical to the model used for comparing the “identical” versus “task-specific” models described above. The inclusion or exclusion of normalized EMG in the stimulus matrix significantly affected model performance regardless of whether phonological features were task-specific (*p*<.001) or identical (*p*<.001). Including information about normalized EMG activity recorded from auxiliary facial electrodes improved model performance (Figure 4A) as shown by the greater number of channels below the unity line. On an individual subject basis, all but two subjects (OP15, OP16) showed a significant difference in model performance across the inclusion or omission of normalized EMG activity as a stimulus feature as assessed by Wilcoxon signed-rank test. When comparing the relative difference between “identical” and “task-specific” models (Figure 3) in the presence or absence of an EMG regressor, models including an EMG regressor showed less of a difference in performance between methods of phonological feature encoding, suggesting that residual EMG decreases the stability of phonological feature tuning across modalities of speech (Figure 4B).

**Figure 4.**
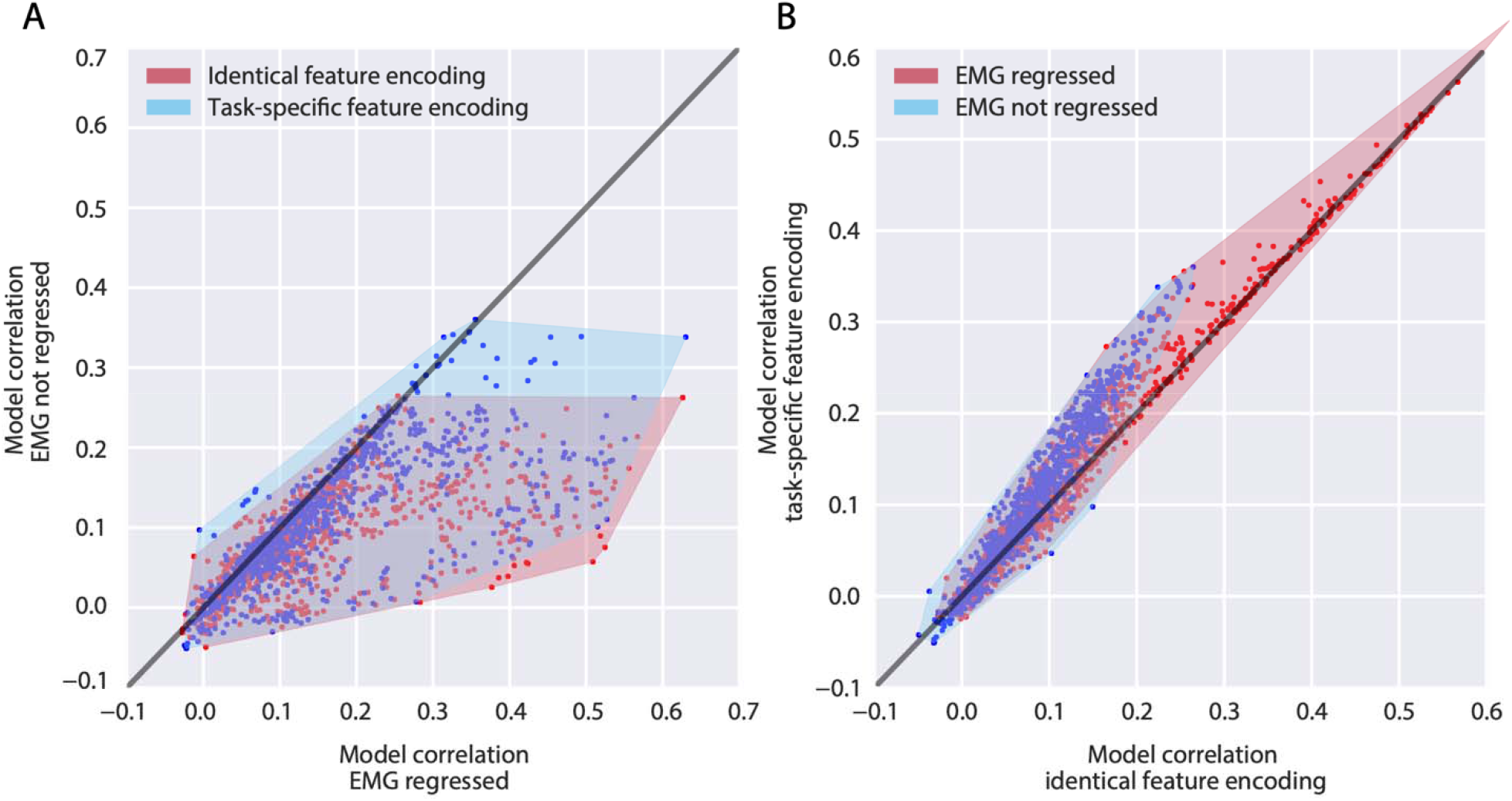
Including EMG as an encoded feature in linear models greatly improves their performance, as well as the stability of phonological feature encoding between perception and production. A: Convex hull plots of individual electrodes’ correlation coefficients with held-out neural response within models that do contain an EMG regressor (x-axis) and those that do not (y-axis), for models that separate phonological feature tuning by task modality (blue) and models that do not (red). Diagonal black line represents unity. B: Convex hull plots of individual electrodes’ correlation coefficients with held-out neural response within models that differentially encode phonological features according to modality of speech (y-axis) and those that do not (x-axis), in the presence (red) or absence (blue) of information about normalized EMG activity recorded from auxiliary facial electrodes. Diagonal black line represents unity. When EMG was regressed, more points lie along the unity line, indicating similar phonological feature tuning.

Trial-specific stimulus features were also ablated to assess their contribution to model performance. Omitting trial modality (i.e., whether a phoneme was produced or perceived) did not significantly affect the linear regression model’s ability to predict the held-out neural response (*p*=0.84). Similarly, ablating information about the predictability of the perception trials did not affect model performance (*p*=0.9). If the EMG regressor is removed in conjunction with trial-specific features, the differences in model performance when trial modality is included or ablated are less profound but still nonsignificant (*p*=0.62). When ablating trial predictability, no changes are observed in significance between inclusion (*p*=0.9) and omission (*p*=0.88) of an EMG regressor, which is expected considering differences in trial predictability are constrained to perception trials where EMG associated with articulation is absent from the response. The ablation of predictability contrast not affecting model performance is in line with the ERP results presented above (Figure 3B). However, ablating trial modality (i.e., perception versus production) not affecting model performance is incongruent with the ERP results, for which a stark contrast between perception and production were observed. The difference in time frame between the ERP analysis (sentence level) and the linear encoding model analysis (phoneme level) may explain the difference between the ERP and linear encoding model results. In other words, sentence-level processing of speech perception and production may involve different neural mechanisms, but at an individual phoneme level, the mechanisms are shared between perception and production. Alternatively, the incorporation of the EMG regressor may be delineating perception and production in the model, making explicit information about trial modality, effectively marking the explicit inclusion of trial type in the stimulus matrix redundant. This explanation is supported by the observation that omission of an EMG regressor substantially impacted model performance.

Taken together, the linear encoding model results suggest that the degree of EMG activity recorded from auxiliary electrodes during the task is an informative characteristic of the stimulus in the context of modeling neural responses to speech. On the other hand, including information about trial type (perception vs. production, predictable vs. unpredictable) was less informative when EMG was included as a regressor. Phonological feature tuning changes across modality of speech, while observed, were reduced with the inclusion of an EMG regressor, which suggests residual EMG artifact in the post-processed signal is responsible for these changes in phonological feature tuning.

## Discussion

### Summary

The results presented in this study demonstrate a difference in EEG responses to perceiving and producing naturalistic stimuli. At the sentence level, a suppression of early auditory components N100 and P200 was observed in speech production relative to perception. These findings are in line with previous literature on speaker-induced suppression and auditory processing more generally. The N100 (and its MEG equivalent N100m) have been theorized as a neural indicator of the efference copy and its suppression has been demonstrated for internally-generated speech compared with externally-generated speech (Martikainen, Kaneko, and Hari 2005; Behroozmand and Larson 2011). The P200 is less directly associated with SIS, with limited studies linking it directly to feedback perturbation (Behroozmand and Larson 2011; Brumberg and Pitt 2019), but it is commonly paired with the N100 in speech perception studies to form the N1-P2 complex (Lightfoot 2016). However, the suppression observed in this study was not intrinsically due to the feedforward nature of speech production, as differences between EEG responses to predictable (feedforward) and unpredictable speech perception were minor.

To investigate what specifically was driving the changes in neural responses to these two modes of speech, we fit linear encoding models describing neural activity as a function of different stimulus features. These features allowed us to test different hypotheses about changes in phonological tuning at the individual feature level versus overall baseline changes during perception and production. Performance of these models were evaluated by how well the weights of the models correlated with held-out EEG response. Including information about auxiliary EMG activity and differentially encoding phonological features during perception and production in the stimulus matrix yielded higher model performance, which suggests these characteristics were represented in the recorded response and offer potential explanations for the suppression of neural activity during speech production relative to perception. However, residual electromyographic artifact may affect interpretability of this approach, considering the inclusion or omission of normalized EMG recorded from facial electrodes substantially affected model performance, such that models that do regress EMG perform better than those that do not. Taken together, these two analyses extend our understanding of speaker-induced suppression by scaling the phenomenon into a more naturalistic context and investigating what specific components of the neural response are being suppressed.

### Potential mechanisms of speaker induced suppression

Previous literature comparing speech production and suppression has identified a neurophysiological effect dubbed speaker-induced-suppression, where internally produced stimuli generate less of a change in neural activity than externally produced stimuli. This study sought to replicate this effect in a more naturalistic setting, as many studies of SIS use low-level acoustic stimuli such as pure tones (Martikainen, Kaneko, and Hari 2005) and single vowels (Niziolek, Nagarajan, and Houde 2013; Houde et al. 2002; Heinks-Maldonado, Nagarajan, and Houde 2006), while many neurolinguistic studies are beginning to prioritize more naturalistic stimuli such as podcasts (Huth et al. 2016; Goldstein et al. 2022), audiobooks (Herff et al. 2015), and movie trailers (Desai et al. 2021) in an effort to better capture how speech and language are used in daily life (Hamilton and Huth 2020). We were able to demonstrate SIS at the sentence level, which is comparatively much more naturalistic than the lower-level characteristics of speech used in previous studies of SIS.

An unanswered question in previous SIS literature is *what* is being suppressed. The observation of speaker-induced suppression in the event-related potential results, combined with a lack of predictability-related suppression in the ERP results and no clear stimulus characteristic contributing substantially to encoding model performance (besides residual EMG artifact), means the question of “*what*” is still very much an open question. One clear difference between speech production and perception is the generation of the efference copy, a feedforward expectation about the content of the upcoming auditory stimulus which is only generated by internally produced speech. A simple explanation is the difference in neural activity between these conditions is due to the presence or absence of the efference copy. The N100, which has previously been identified as a biomarker of the efference copy (Brumberg and Pitt 2019), was suppressed during speech production in our study. To expand on the N100 suppression observed in the ERP, we ablated specific stimulus characteristics from linear encoding models and observed how the absence of a specific aspect of the stimulus affected the model’s ability to predict the neural response. We observed differential phonological feature encoding across perception and production using this approach, which offers a potential explanation for the content of the efference copy (if we believe that is what is suppressed during SIS). However, the observation that regressing EMG activity increased stability of phonological feature tuning across modalities, coupled with the ablation of information about stimulus modality, suggests that residual EMG artifact may be driving the differences between phonological feature tuning during perception and production (Figure 4). Additionally, the structure of the receptive fields themselves was relatively consistent – that is, the brain does not shift towards representing different phonological features during perception and production (Figure 3A).

The alternative conclusion to draw is that speech perception involves additional processing costs not necessary during speech production, such as the segmentation into invariant linguistic representations (e.g., phonological features (Mesgarani et al. 2014)) from a continuous acoustic stimulus. During speech production, linguistic representations are used during pre-articulatory planning, meaning the invariant representations are already available to the language network, negating the need for additional processing costs associated with segmentation. Other information encoded in neural responses to speech perception may also be absent during speech production, which would help explain what is being suppressed during SIS. For example, onset and sustained response profiles have been observed in superior posterior temporal gyrus (Hamilton, Edwards, and Chang 2018). If one of these response types is redundant with information contained in the efference copy, it may not be present in responses to internally generated speech. Future studies will determine if this is indeed the case.

### Pre-articulatory Readiness Potential in Speech Production

In our event-related potential analysis, we observed a positive deflection in the grand average ERP (Figure 2B) that began ∼200ms before articulation and peaked ∼100ms before articulation present in the speech production trials. We believe this activity to be related to feedforward linguistic and motoric preparation that must take place prior to articulation. Before articulation, a communicative desire must be morphologically, syntactically, and lexically encoded before it is transformed into a motor program for the speech articulators (Levelt 1993; Flinker et al. 2015; Tourville and Guenther 2011). The exact pre-articulatory stages of speech production are difficult to dissociate with this task, as there is no epoched timing information available as to when these processes occur in a naturalistic context; however, the presence of this pre-articulatory activity exclusively during speech production motivates these stages as an explanation. Pre-stimulus activity was also observed in the grand average during perception trials in the form of positive activity starting at -600ms and peaking at stimulus onset. This activity may be related to predictive components of speech perception, as feedforward processing is an important aspect of successful speech perception (Poeppel and Monahan 2011; Hamilton et al. 2021; Heald and Nusbaum 2014). This speculation is supported by the structure of the task allowing participants to anticipate when they would hear a sentence; however, this task was not operationalized in a way that allows a more granular analysis of this phenomenon. Notably, for both perception and production, the polarity of the pre-stimulus activity was inconsistent from subject-to-subject. This internal inconsistency suggests the activity is not related to previously described ERP components (e.g., readiness potential/Bereitschaftspotential) as these components have a canonical negative polarity (Yoshida et al. 1999; Wohlert 1993; Jahanshahi and Hallett 2003).

### Influence of EMG Artifact

In any noninvasive neuroimaging study of speech production, movement artifacts caused by articulation are a concern to the integrity of the data. Traditionally, EEG analyses of speech production have sidestepped addressing EMG by requesting participants “imagine” speech while not moving the articulators or shifting the analysis window to a time outside when articulatory movement is occurring. However, there have been recent successful attempts at directly analyzing the window of overt articulation up to the phrase level (Ries et al. 2021). In this experiment, stimuli consisted of four-word tongue twisters. We extend these results by scaling up to the sentence level with evoked responses to speech appearing relatively cleaned of EMG artifact as evidenced by the integrity of the N100 and P200 components. A reason to assume that residual EMG is affecting the results is the differing performance of encoding models that do or do not regress EMG (Figure 4). Models that accounted for EMG as a stimulus characteristic on the whole outperformed models that did not, which means there is variance remaining in the post-processed data that is well explained by electromyographic activity. The inclusion of an EMG regressor was only made possible by recording facial muscle activity using auxiliary electrodes in conjunction with the EEG, akin to how EEG researchers will record auxiliary vertical and horizontal EOG to assist with artifact correction. While previous research has demonstrated blind source separation-based artifact correction techniques are sufficient in correcting EMG artifact for event-related potential analysis, the substantial difference in model performance when this normalized EMG activity was ablated from the stimulus matrix leads us to strongly recommend the use of auxiliary EMG recordings to any researchers who wish to fit similar linear encoding models to speech production data. Furthermore, we only recorded single-channel EMG, while there are a plethora of facial muscles that contribute to EMG artifact in the electroencephalogram. It is possible that including activity from multiple auxiliary channels as a regressor in linear encoding models would further improve their performance, but future research is needed to substantiate this claim.

There are several reasons we do not believe the residual EMG in our response nullifies the interpretation of this study’s results. First, the integrity of purely auditory responses is preserved after post-processing as evidenced by evoked responses to inter-trial click tones, which suggests the evoked responses seen at the sentence level are not false positives caused by EMG artifact. Second, despite the contribution of EMG to linear encoding models, we observe strong phonological feature tuning consistent with previous research (Hamilton et al. 2021; Desai et al. 2021). Third, including EMG as a regressor in linear encoding models ensures that phonological feature tuning (or a similar feature space of interest) is not obscured or affected by muscle artifact. Lastly, evoked responses to sentence onset contained robust N100 and P200 components that would not be visible in the presence of substantial noise from EMG.

### Stimulus Predictability

A manipulation of predictability was included in the present study to assess the hypothesis that speaker-induced suppression is associated with general feedforward auditory processing and not an intrinsic characteristic of corollary discharge during speech production. Differences between predictable and unpredictable perceptual trials were small, with only three individual subjects demonstrating a significant difference. Predictability-related suppression is well-supported by the literature, but appears to be a separate mechanism from SIS (Lester-Smith et al. 2020; Goregliad Fjaellingsdal et al. 2020; Bendixen et al. 2014; Astheimer and Sanders 2011). One of the most well-documented instances of predictability-related suppression is mismatch negativity (Hawco et al. 2009; Näätänen et al. 2007), which is commonly studied using an oddball stimulus paradigm. Our study presented the predictable and unpredictable perceptual trials in blocks of 50 trials each, which means participants could identify when perceptual stimuli would be unpredictable, a fundamental difference from the oddball tasks where deviant stimuli are presented randomly. We chose not to present unpredictable stimuli in an oddball fashion because our perceptual stimuli were generated from the recorded productions of the participant. Thus, to generate the full range of unpredictable perceptual stimuli in our task, a full block of production trials is needed, and collecting this as a baseline before introducing oddball unpredictable stimuli would greatly extend the time of our recording sessions, and we judged more repetitions of each condition to be more important to our research questions. The block design of the task may also cause listeners to adapt to the randomly shuffled perceptual stimuli over the course of the block. In feedback perturbation studies, it has been demonstrated that listeners are more likely to correct to the altered feedback if the perturbations are presented in predictable on-off blocks compared to at random (Lester-Smith et al. 2020).

Beyond predictability, several papers on SIS have posited active versus passive listening as the cognitive difference between speaking and listening (Houde et al. 2002; Brumberg and Pitt 2019). Speech production requires active listening to monitor for potential errors in the speech signal which can subsequently be corrected, while speech perception does not. Many psycholinguistic studies have required active listening of their participants by having them attend to specific components of the stimulus which the participants must answer questions about later in the study. An exploration of differences in the N100 between active and passive speech perception trials could potentially provide further information on the amodal mechanism behind speaker-induced suppression.

### Conclusion

The current study uses naturalistic sentence-level speech perception and production to examine how EEG responses to speech differ across modality and predictability of speech. After correcting for electromyographic artifact in the data via canonical correlation analysis, we demonstrate speaker-induced suppression in production relative to perception as demonstrated by amplitude and latency of the N1-P2 complex epoched to sentence onset. No major findings were observed for predictability contrasts in the perception trials. Next, we examined phonological feature tuning across these behavioral conditions via a series of linear encoding models. A difference in tuning was observed such that separating phonological feature encoding across perception and production improved model performance, but these gains were attenuated by the inclusion of normalized EMG as a regressor, suggesting that residual EMG in the model decreases stability of phonological tuning across modalities of speech. This research hopes to illuminate differences in electrophysiological responses to perception and production, which contributes to the study of disorders in which these mechanisms are disrupted, such as stuttering, schizophrenia, and Parkinson’s disease, and motivate future study of naturalistic speech production with noninvasive electroencephalography.

## Data Availability Statement

Code for reproducing the analyses in this manuscript can be found at https://github.com/HamiltonLabUT/speaker_induced_suppression_EEG/. The EEG dataset and corresponding event files can be downloaded at https://doi.org/10.17605/OSF.IO/FNRD9.

## Author Contribution

Funding for this study was acquired by LSH. GLK and LSH conceptualized the study along with consultation from RLS. GLK and LSH contributed to project administration. GLK supervised AM, NC, JH, CV, VRM, CT, CH, and PP in investigation and data curation. Formal analysis, methodology, software, validation, and visualization were performed by GLK and LSH. GLK prepared the initial draft. All authors contributed to review and editing of the manuscript. The authors have no conflicts of interest or relevant disclosures of competing interests.

## Acknowledgements

The authors would like to thank Maansi Desai, Mary Lowery, and Ian Griffith for their assistance in data collection. The authors additionally would like to thank Stéphanie Riès for her assistance with the preprocessing steps of the experiment.

